# Prevalence of the Calvin-Benson-Bassham cycle in chemolithoautotrophic psychrophiles and the potential for cold-adapted Rubisco

**DOI:** 10.1101/2024.08.01.606197

**Authors:** Kaitlin Harrison, Josephine Z. Rapp, Alexander L. Jaffe, Jody W. Deming, Jodi Young

## Abstract

The act of fixing inorganic carbon into the biosphere is largely facilitated by one enzyme, Rubisco. Beyond well-studied plants and cyanobacteria, many bacteria use Rubisco for chemolithoautotrophy in extreme environments on Earth. Here, we characterized the diversity of chemolithoautotrophic Rubiscos in subzero environments. First, we surveyed subzero environments and found that the Calvin-Benson-Bassham cycle was the most prevalent chemolithoautotrophic pathway. Second, we uncovered potential for chemolithoautotrophy in metagenomes from two distinct subzero, hypersaline Arctic environments: 40-kyr relic marine brines encased within permafrost (cryopeg brines) and first-year sea ice. Again, the Calvin-Benson-Bassham cycle was the dominant chemolithoautotrophic pathway in both environments, though with different Rubisco forms. From cryopeg brine, we reconstructed four metagenome-assembled genomes with the potential for chemolithoautotrophy, of which the sulfur-oxidizing genus *Thiomicrorhabdus* was most abundant. A broader survey of *Thiomicrorhabdus* genomes from diverse environments identified a core complement of three Rubisco forms (II, IAc, IAq) with distinct patterns of gain and loss. We developed a model framework and compared these different Rubisco forms across [CO_2_], [O_2_], and temperature. We found that form II outcompetes form I at low O_2_, but cold temperatures minimize this advantage. However, further inspection of form II from cold environments uncovered signals of thermal adaptation of key amino acids which resulted in a more exposed active site. These modifications suggest that these form II Rubisco proteins may have unique kinetics or thermal stability. This work can help address the limits of autotrophic functionality in extreme environments on Earth and other planetary bodies.

**Importance:** Autotrophy, or the fixation of inorganic carbon to biomass, is a key factor in life’s ability to thrive on Earth. Research on autotrophy has focused on plants and algae, but many bacteria are also autotrophic and can survive and thrive under more extreme conditions. These bacteria are a window to past autotrophy on Earth, as well as potential autotrophy in extreme environments elsewhere in the Universe. Our study focused on dark, cold, saline environments, which are likely to be found on Enceladus and Europa, as well as in the Martian subsurface. We found compelling evidence of cold adaptation in a key autotrophic enzyme, Rubisco, which could expand the known boundaries of autotrophy in rapidly disappearing icy environments on Earth. We also present a novel model framework that can be used to probe the limits of autotrophy not only on Earth but also on key astrobiological targets like Enceladus and Europa.

## Introduction

Autotrophy, or the fixation of inorganic carbon to complex organic molecules, is required for almost every ecosystem on Earth. In addition, the earliest known life on Earth is often believed to be autotrophic [1–3]. While recent work on protein evolution has cast doubt on our understanding of the physiology of the last common ancestor [4], what is not in doubt is the early emergence of autotrophy in the history of life on Earth and its importance in a thriving Earth biosphere.

There are seven naturally occurring carbon fixation pathways found on Earth [5,6]. However, estimates of global carbon fixation suggest that the Calvin-Benson-Bassham (CBB) cycle is responsible for >99% of the autotrophy on the planet [7]. This cycle is the only carbon fixation pathway in oxygenic photosynthesis and thus is found in every plant, cyanobacterium, and alga; however, the CBB cycle is also utilized in chemolithoautotrophs, coupled to a diverse set of prokaryotic metabolisms including oxidation of sulfur, iron, ammonia, and nitrite [7]. While likely not the earliest carbon fixation pathway to evolve [3], it is found in almost every habitat on the planet regardless of temperature, pressure, salinity, and oxygen concentration. Due to its prevalence, enzyme diversity in the CBB cycle is much better sampled than in any of the other autotrophic carbon fixation pathways, making it an incredibly useful tool to study the adaptation of carbon fixation to various environmental conditions.

The maximum speed of the CBB cycle is typically controlled by the enzyme ribulose 1,5-bisphosphate carboxylase/oxygenase (EC 4.1.1.39), also known as Rubisco. In the CBB cycle, Rubisco catalyzes the step in which CO_2_ is added to a 5-carbon organic sugar (ribulose 1,5-bisphosphate, or Rubp) to form two 3-phosphoglycerate molecules. Thus, Rubisco is directly responsible for the conversion of inorganic carbon to organic carbon. Despite its critical role in the CBB cycle, Rubisco is not well adapted to our current atmospheric CO_2_ and O_2_ concentrations [8]. Rubisco is competitively inhibited by oxygen, resulting in the production of 2-phosphoglycolate (2-PG), a toxic molecule whose removal results in the loss of recently fixed carbon. In plants (where this process is called photorespiration), up to 49% of gross primary production can be lost to this phosphoglycolate salvage (PGS) pathway [9]. During the Archean, when Rubisco evolved [10], this side reaction would have been rare due to the relatively high CO_2_ levels and extremely low O_2_ levels. However, as O_2_ levels have risen and CO_2_ levels have dropped, PGS has become a significant source of carbon loss. Rubisco is also poorly suited for the temperatures at which it typically operates across most of Earth’s surface. In vitro assays demonstrate that Rubisco has maximum activity between 45 and 60°C [11], temperatures that most of the ocean and land never reach. At Earth’s average surface temperature (15°C), Rubisco has somewhere between 4 and 13% of its peak activity [11].

Most of the research on Rubisco to date has focused on Rubisco form IB, found in plants, some cyanobacteria, and green algae. This form represents only a small amount of the diversity of Rubisco in current organisms. There are four known forms of Rubisco, which are often distinguished further by sub-type and are present in different branches of life [12]. Form I Rubisco is a hexadecamer, made of 8 large subunits and 8 small subunits. Form I can be further split into subtypes (forms IA–IE). Form I subtypes, excluding form IB, are present in various algae and bacteria. Form II Rubisco, which is only composed of a single large subunit, can form a wide range of oligomers depending on species; it is found in bacteria and some algae (dinoflagellates). Form III Rubisco and form IV Rubisco are typically not thought to be involved in autotrophic carbon fixation, although there are exceptions [5]; typically, form III Rubisco is responsible for nucleoside salvage in prokaryotes, especially archaea, while form IV does not carboxylate RuBP and is thought to be involved in a range of different processes [12]. There are also a number of other forms that have been discovered more recently, including I’, Iα, and II/III [12]. (For a more thorough review of Rubisco’s diversity and evolutionary history, see [12].) While many organisms, including plants, only contain one form of Rubisco, many proteobacteria that utilize the CBB cycle often encode multiple forms of Rubisco [13].

Despite decades of Rubisco research, there remain outstanding questions about why this tremendous amount of enzyme diversity exists. One hypothesis is that diversity is driven by the environmental pressures of the aforementioned mismatch between optimal conditions for Rubisco activity and the current conditions on Earth. The ability of Rubisco to adapt to a particular environment remains debated due to its extremely slow rate of evolution [14]. However, the currently available data are dominated overwhelmingly by plant Rubisco form IB [11] measured at a canonical 25°C [15] and thus lack power for testing environmental adaptation. Many non-IB forms of Rubisco have kinetics that vary significantly from form IB [16,17]. Most studies have focused on a suspected trade-off between carboxylation speed and affinities for CO_2_ over O_2_ [16,17]. An increase in speed is advantageous under high CO_2_ [8], and many organisms have evolved an intracellular carbon concentrating mechanism to elevate CO_2_ around Rubisco [18]. One such carbon concentrating mechanism is the carboxysome, found in cyanobacteria and some chemolithoautotrophic bacteria. This bacterial protein structure excludes O_2_ while allowing CO_2_ to be concentrated around Rubisco [18].

Despite all of this diversity, little is known about the potential temperature sensitivities of the various Rubisco forms. In form IB, decreasing temperature increases the CO_2_ specificity (S_C/O_) and the CO_2_ affinity (K_C_) while carboxylation rate (*k*_cat,C_) decreases [19]. There are also minor differences in temperature sensitivity between plants typically found in cold versus warm environments [19]. Few measurements of temperature dependence have been made outside the form IB Rubiscos [19]. Even fewer are from extremophiles, and those that do exist are more commonly from thermophiles than psychrophiles [19]. The scant measurements available from psychrophiles suggest opposite results: a form ID from a psychrophilic alga (a diatom) has temperature kinetics similar to mesophilic diatoms [20], while form ID Rubiscos from cold-temperature seaweeds have higher *k*_cat,C_ at cold temperatures than their mesophile counterparts [21]. More research is needed to clarify the effects of temperature on the various forms of Rubisco, especially those from extreme environments.

Subzero environments are extreme environments facing significant changes in the next decades due to climate change. The Arctic is predicted to be ice-free in the summer by the 2030s [22], and the infiltration of permafrost melt into homes and food storage is a potential hazard to residents of the North, especially Indigenous communities [23]. Taxa-specific traits are needed to model the polar carbon fixation response to these changes [24]. Beyond Earth, the 2020 Astronomy Decadal Survey has called for more research on poly-extreme environments as analogues for similar environments on exoplanets [25]. Permafrost brines and sea ice are particularly analogous to conditions on the icy moons of Jupiter and Saturn, especially Europa and Enceladus, which are considered high priority targets in the search for life in our solar system [25].

Here, we reviewed autotrophic pathways and taxa in subzero environments. Then, we quantified the abundance of Rubisco forms found within metagenomes from ancient marine brines within the permafrost (cryopeg brines) and sea ice. We further explored Rubisco diversity in an autotrophic bacterial genus, *Thiomicrorhabdus*, due to its prevalence in subzero environments. *Thiomicrorhabdus* genomes contain multiple Rubisco genes and unique amino acid substitutions that hint at adaptation of Rubisco to different environmental stressors, particularly subzero temperatures. Finally, we developed a theoretical model to examine the tradeoff between Rubisco forms under different CO_2_, O_2_, and temperature regimes using currently available data but presenting a framework that can be utilized broadly.

## Results

### Autotrophy potential at subzero temperatures

A survey of the current literature revealed that five of the seven known natural carbon fixation pathways are present in chemoautotrophs at subzero conditions (Table 1). The two missing pathways are the 4-hydroxybutyrate/dicarboxylate (4-HB/DC) cycle, only found in archaea [5], and the reductive glycine pathway, only recently discovered [6]. The subzero environments surveyed included high-latitude seawater in winter, terrestrial brines, endolithic habitats, polar brines, and sea ice. While not an exhaustive list, we found that across these diverse environments, the CBB cycle was present in every environment where a carbon fixation pathway was detected, regardless of temperature variability or oxygen concentrations. Brines at Lost Hammer Spring in the Arctic and within rocks at Blood Falls, Antarctica, contained organisms with the Wood-Ljungdahl pathway and the reductive tricarboxylic acid (rTCA) cycle in addition to the CBB cycle. The rTCA cycle was also present in brines at Gypsum Hill Spring in the Arctic. Seawater under the Ross Sea ice shelf in Antarctica had organisms with the widest variety of chemoautotrophic pathways, with the CBB cycle, the rTCA cycle, the 3-hydroxyproprionate (3-HP) bi-cycle, and the 3-hydroxyproprionate/4-hydroxybutyrate (3-HP/4-HB) cycle all found there. Archaea with an autotrophic pathway were only detected under the Ross Sea ice shelf.

**Table 1:**
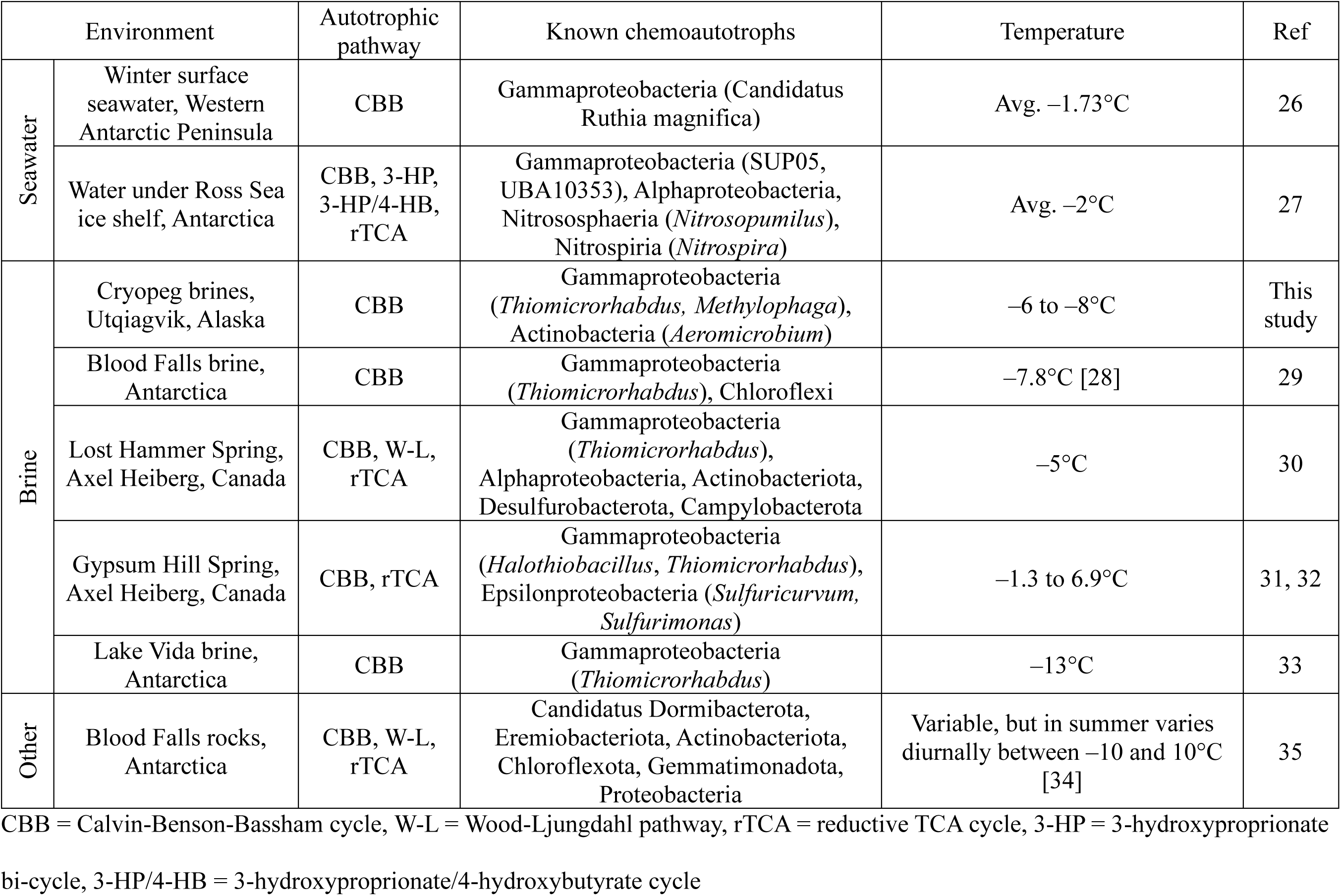
Survey of potential chemoautotrophy in subzero environments.

Proteobacteria were common chemolithoautotrophic members of the community, especially Alpha- and Gammaproteobacteria. One of the few taxa reported to the genus level was *Thiomicrorhabdus*, which was observed in four different high salinity habitats across both poles.

### Rubisco forms within cryopeg brines and sea ice

To take a closer look at the CBB cycle in subzero environments, we searched for Rubisco sequences found within metagenomes from Arctic cryopeg brines and sea-ice brines and cores that we had collected previously from Utqiaġvik in 2017 and 2018 (for full description of sampling see [36,37]). Sea-ice brines had an average temperature and salinity of –3.5°C and 77 ppt. Sea-ice core melts had an average temperature and bulk salinity of –4°C and 18 ppt. Cryopeg brines were considered anoxic *in situ*, with an average temperature and salinity of –6°C and 123 ppt.

A total of 774 redundant sequences attributed to the Rubisco large subunit were detected across all samples, with the vast majority (693 sequences or 89.5%) derived from sea ice. On average, scaffolds containing a Rubisco sequence were responsible for only a small fraction (approximately 0.001%) of mapped reads from each sample. Each sequence was assigned to a Rubisco form, and most of the major forms of Rubisco were detected in both cryopeg brine and sea-ice samples, with the notable exception of form III. None of the recently discovered forms (i.e. I’, Iα, II/III) were detected.

Rubisco sequences from sea-ice samples were predominately of form ID (76% of mapped reads to Rubisco scaffolds) with a lesser fraction of form IB (16%), all attributed to a variety of diatoms and green algae. As these are photosynthetic organisms and not the target of this study, they were removed from further analysis. The remaining 8% were of bacterial origin, mostly attributed to Gamma- and Alphaproteobacteria. These sequences were from 6 distinct Rubisco groupings: forms IA and IC, form II, and three different form IV clades. Fig. 1 shows the proportion of these different bacterial forms in both cryopeg brines and sea-ice samples.

**Fig. 1:**
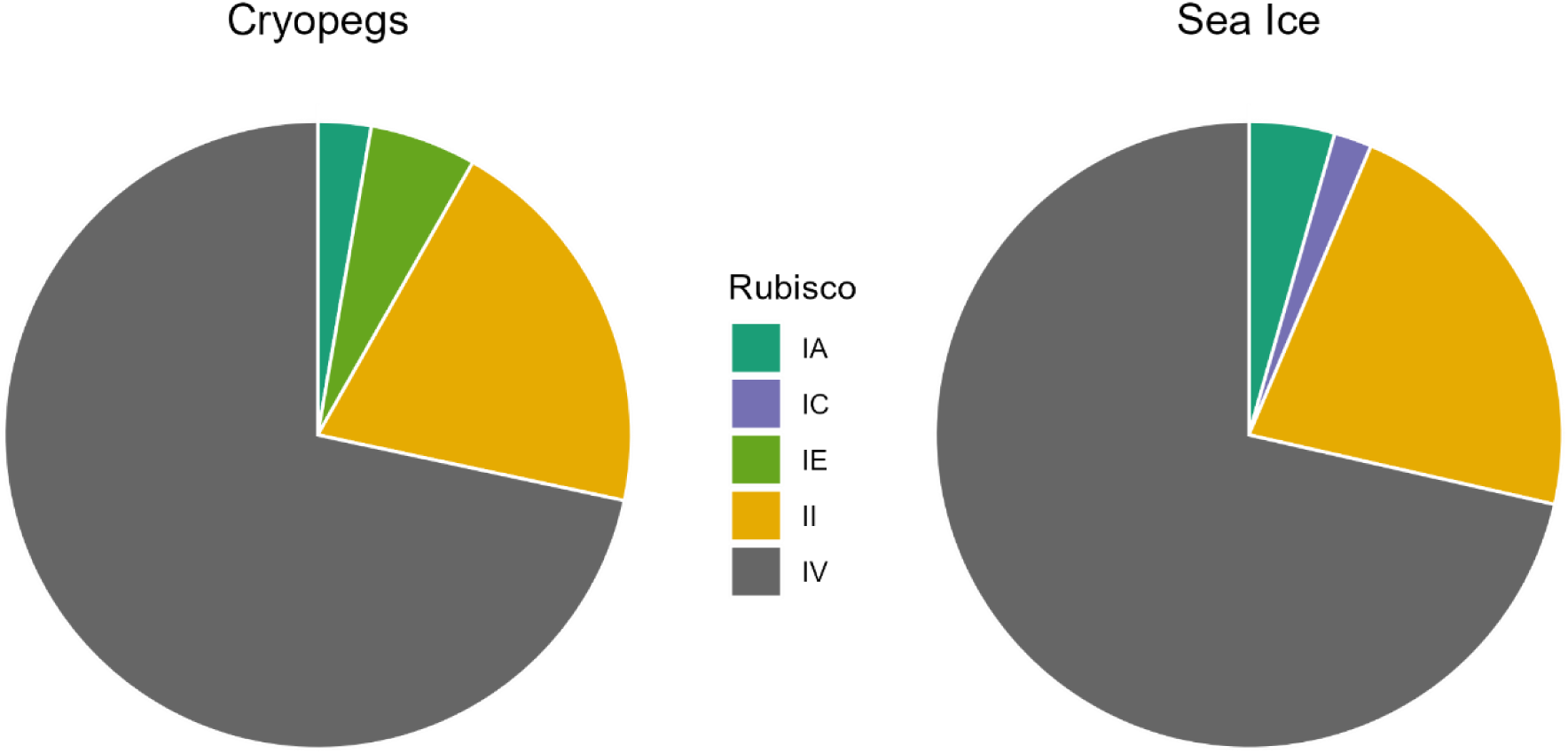
Rubisco forms in metagenomes from cryopeg brines and sea ice. Proportion of bacterial Rubisco abundance attributed to each form within cryopeg brine and sea ice metagenomes. Abundance calculated from mapped reads to Rubisco-encoding scaffolds. No Form III was detected in either sea ice or cryopeg brine samples.

Within the cryopeg brines, autotrophic Rubisco sequences (i.e. forms I and II) were found in 5 of the 6 samples, with the outlier having a bacterial community and geochemistry distinct from the others [37]. All of the Rubisco sequences were of bacterial origin. No autotrophic eukaryotes or archaea were present in the cryopeg brines. Most of the Rubisco sequences were either form IV (72% of mapped reads) or form II (20%), and the remainder was composed of forms IA and IE.

In both sea-ice samples and cryopeg brines, form II Rubisco was more abundant than form IA (5.0x and 7.3x more mapped reads respectively). Half of the form II and all of the form IA Rubisco sequences in the cryopeg brines were attributed to *Thiomicrorhabdus*, a genus of Gammaproteobacteria, which is known to have multiple forms of Rubisco [38].

### Autotrophic MAGs within cryopeg brines

Of the metagenome-assembled genomes (MAGs) reconstructed from Utqiaġvik cryopeg (UC) brines, five were identified that encoded a full autotrophic pathway. Three MAGs belonged to Gammaproteobacteria (two different *Thiomicrorhabdus* MAGs, hereafter called UC1 and UC2, and a *Methylophaga* MAG) and one Actinobacteria MAG (*Aeromicrobium*), all with a full CBB cycle (Table 2). The fifth MAG was identified as belonging to the genus *Sulfurospirillum* and encoded the rTCA cycle. As the genus *Sulfurospirillum* has never been shown to grow autotrophically, despite the presence of the full rTCA cycle [39,40], it was not considered autotrophic for this analysis. Full details of MAG completeness and contamination are shown in Table 2. The contribution of putative autotrophic members to the total cryopeg brine community was small, with 0.5–1.5% of the total reads mapping to the autotrophic MAGs across all samples (Fig. S1).

**Table 2:**
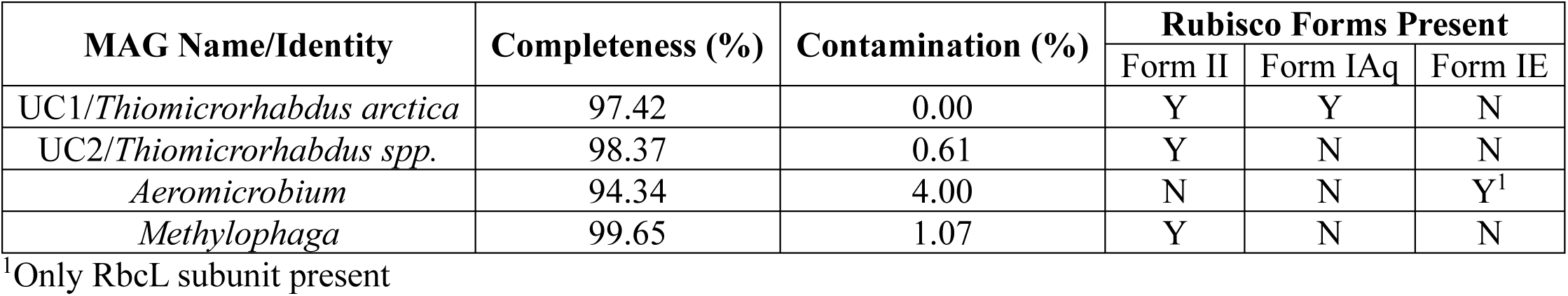
Cryopeg brine MAG completeness and Rubisco complement.

Average nucleotide identities (ANI) of the cryopeg *Thiomicrorhabdus* MAGs UC1 and UC2 (Fig. S2) identified UC1 as a new strain of *T. arctica* (98.62% ANI), while UC2 was most likely a new species, with the closest match to *T.* sp. NP51 (89.79% ANI), a strain of *Thiomicrorhabdus* isolated from a Canadian cold saline spring [41].

The UC1 genome contained both a form II and a form IA Rubisco, consistent with the previously isolated and sequenced strain of *T. arctica* [38]. UC2 only contained a form II Rubisco. The neighboring genes and their order around the form II cassette remained the same as those surrounding the form II and form IA cassette in *T. arctica*, the closest cultured relative of UC2 (Fig. S3), leading us to conclude that this was a true loss of form IA from UC2.

### *Rubisco in* Thiomicrorhabdus

As *Thiomicrorhabdus* was present in our cryopeg brine samples and is known to be actively fixing carbon in subzero environments [29,41], we next focused on Rubisco in *Thiomicrorhabdus*. We compared the recovered cryopeg sequences to other *Thiomicrorhabdus* Rubisco sequences from diverse habitats to infer whether specific Rubisco forms are preferentially retained or lost under subzero conditions.

There are 45 publicly available *Thiomicrorhabdus* genomes (both MAGs and cultured isolates) from a variety of habitats that encompass a wide range of temperature and salinity, with cultured strains able to grow from –2 to 45°C and 0 to 1700 mM NaCl (approximately 0 to 99 ppt) (Table S1). Half of the genomes with known sampling locations were found at hydrothermal vents. Two genomes were sampled from subzero environments, and an additional seven cultured isolates grow below 5°C. Known sampling locations are displayed in Fig. S4. Where available, the temperature and salinity information for each genome, as well as quality and source information, are given in Table S1. Of the 45 genomes, 22 passed quality control (<5% contamination, >75% completeness, and de-replication) for our analysis; 17 of these were genomes from cultured isolates.

A maximum likelihood phylogenetic tree was constructed from these 22 publicly available *Thiomicrorhabdus* genomes, and 2 genomes from a closely related genus, *Hydrogenovibrio,* as an outgroup (Fig. 2). Our cryopeg MAGs UC1 and UC2 and other genomes isolated from cold environments formed a distinct clade (Clade 2) based on ANI. Rubisco forms found in each genome are displayed alongside. All *Thiomicrorhabdus* genomes encoded at least one Rubisco gene with many encoding up to 3 different forms: form II and two forms of IA, one ‘free’ (IAq), and the other carboxysome-associated (IAc) (Fig. 2B). Both form II and form IAq were found in a cassette with their associated transcriptional regulator LysR gene and Rubisco activase (CbbQ/CbbO dimer pair) (Fig. S6). Form IAc was surrounded by carboxysome genes, as seen in Scott et al. 2019 [47]. Rubisco gene duplication was only detected in *T. sediminis*, which contained two copies of form IAc. Both *Hydrogenovibrio* genomes encoded all three different forms of Rubisco found in the *Thiomicrorhabdus* genomes.

**Fig. 2:**
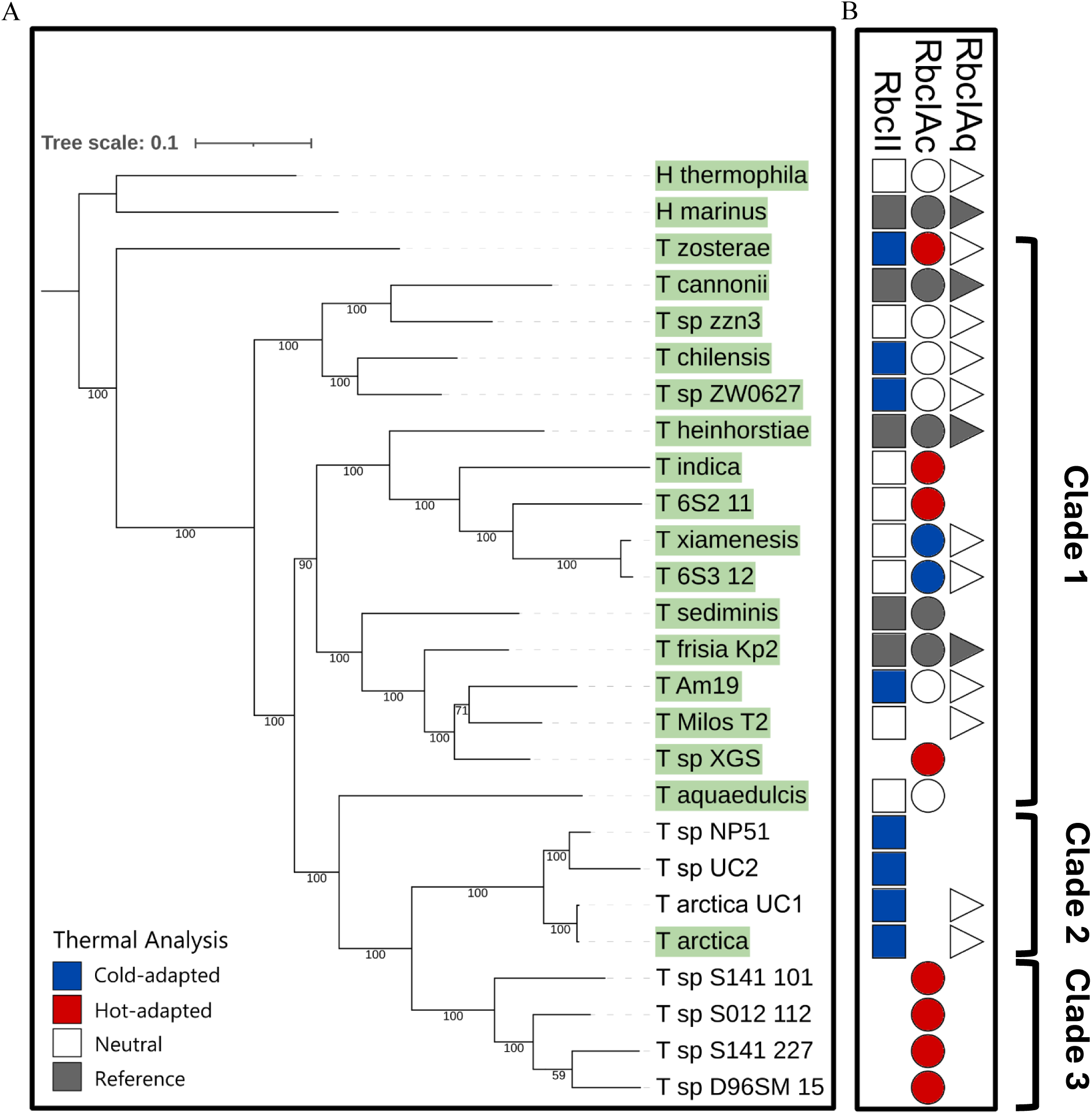
Phylogenetic tree of *Thiomicrorhabdus* genomes alongside Rubisco gene complement and results from amino acid thermal analysis. A) Maximum-likelihood phylogenetic tree of 22 *Thiomicrorhabdus* genomes created by IQTREE2 [42] using the Bacteria 71 gene collection [43–45]. *H. marinus* and *H. thermophila* designated as outgroup. Bootstrap values for each node are depicted on the branch. Cultured isolate genomes highlighted in green. *T. arctica* UC1 and *T.* sp. UC2 were placed on the tree manually due to lack of ribosomal genes, using ANI as a proxy. B) Presence/absence of Rubisco forms and result of thermal adaptation. Shape represents presence of form: square = form II, circle = form IAc, triangle = form IAq. Color indicates result of thermal analysis: blue = cold-adapted, red = hot-adapted, outline-only = neutral, grey = used as reference for thermal analysis. Tree and datasets visualized with iTOL [46]. See Fig. S5 for an expanded Clade 3 that includes two other genomes in Clade 3 that passed the quality control but did not contain a full Rubisco gene.

The distribution pattern of Rubisco forms within *Thiomicrorhabdus* is complex and cannot be explained by a few instances of gene loss or gain. Interestingly, all genomes isolated from subzero environments encoded a form II but no form IAc (Fig. 2, clade 2), whereas a clade of genomes from hydrothermal vents only encoded form IAc (Fig. 2, clade 3). We acknowledge that the absence of a gene is difficult to confirm in a MAG; however, 5 of the MAGs are ≥90% complete, including all of the Clade 2 MAGs and two of the Clade 3 MAGs, and the same complex distribution of Rubisco genes remained when only considering genomes from cultured isolates. For example, within Clade 1, cultured isolates *T. immobilis* (Am19), *T. sp.* Milos-T2, and *T. sp.* XGS-01 are all closely related based on ribosomal genes but encode different complements of Rubisco genes.

### Theoretical framework to compare Rubisco kinetics under diverse environmental conditions

It has been hypothesized that multiple Rubisco forms within proteobacteria are advantageous under different environmental conditions [13,47]. To test this hypothesis in *Thiomicrorhabdus*, we derived a theoretical framework and compared different Rubisco forms within a matrix of CO_2_, O_2_, and temperature conditions. As kinetics data from *Thiomicrorhabdus* is unavailable, we used the published kinetics data of the three different Rubisco forms (form II, form IAq, and form IAc) present in *Hydrogenovibrio marinus* [48,49]. For each form, carboxylation speed (*k*_cat,C_), half-saturation constant for CO_2_ (K_C_) and O_2_ (K_O_), and CO_2_/O_2_ specificity (S_C/O_) were modeled across CO_2_ and O_2_ concentrations (see Methods and Table S2). The influence of temperature on each kinetic parameter was also modeled for 0, 25, and 55°C, based on trends from Galmés et al. 2016 [19]; however, due to the limited available data, the temperature dependence equations of the form IAc and IAq Rubisco kinetics were very similar, especially for *k*_cat,C_ (Fig. S7). For full details see Methods and Tables S2–S5.

The carboxylation rate over the CO_2_/O_2_ parameter space at the three temperatures was calculated for the three different Rubisco forms, considering just Rubisco (gross), or including the loss of CO_2_ due to PGS using the canonical plant pathway or an alternate bacterial pathway (Fig. 3, Table S5). All three Rubisco forms showed the same basic pattern: the fastest rate is reached under low O_2_, high CO_2_, and high temperature. When not accounting for carbon loss through the PGS pathway, form II was always faster than form IAc and IAq (Fig. 3A,D,G). However, incorporation of the PGS pathway resulted in form IAc outcompeting form II under warm, oxygenated conditions with low CO_2_ (Fig. 3B,E,H). Form IAc outcompeted form II to a greater extent when using the alternate bacterial pathway for PGS (Fig. 3C,F,I). The form IAq enzyme was outcompeted by IAc and II under all conditions.

**Fig. 3:**
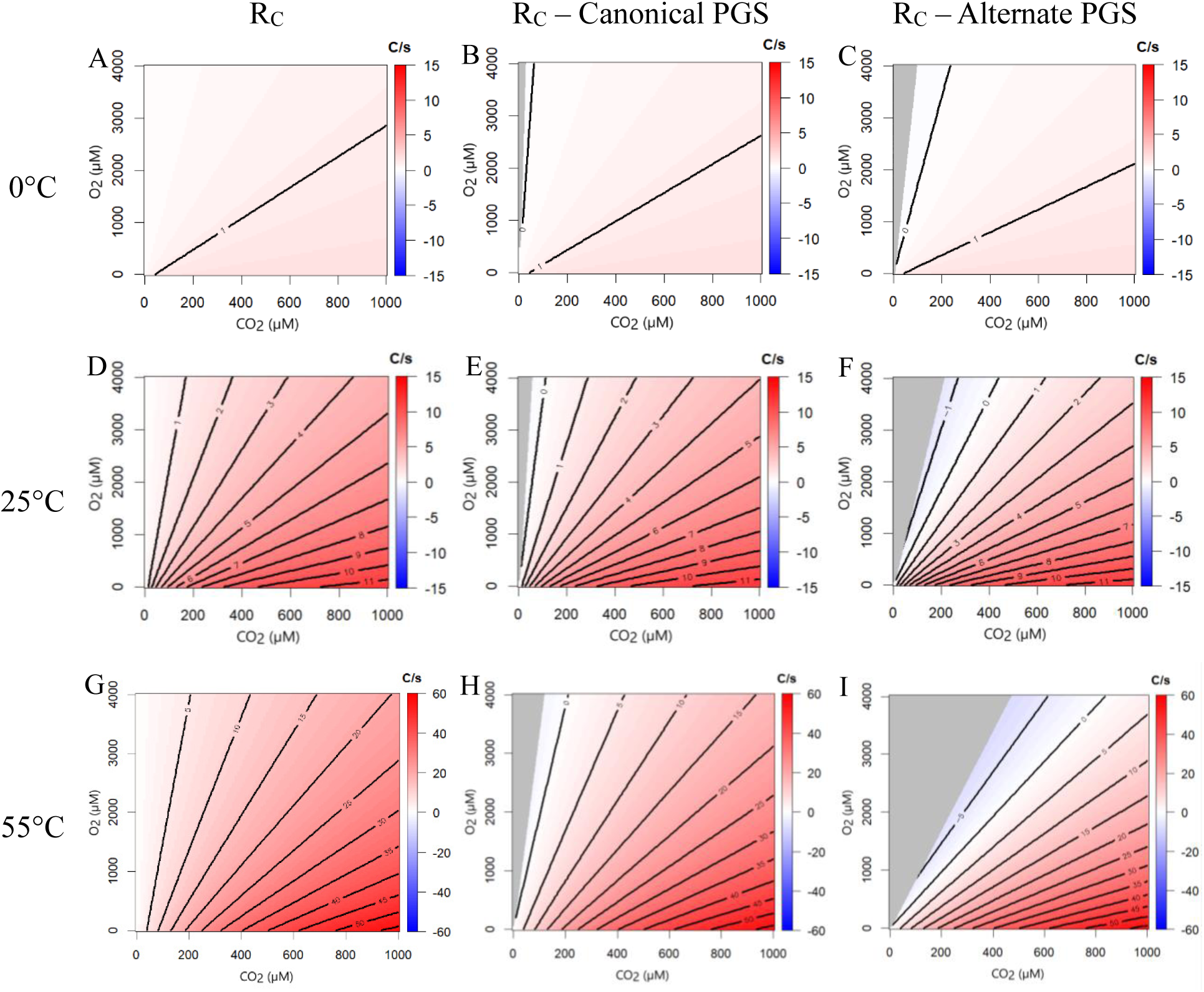
Theoretical kinetics model of Rubisco net carboxylation. Modeled Rubisco carboxylation speed for forms II – IAc at varying CO_2_ concentration, O_2_ concentration, and temperature. Red indicates form II > IAc, blue indicates form IAc > II, with shading indicating magnitude in carbons per second (C/s). Gray shading at low CO_2_ indicates conditions where there was a net loss of carbon. A, D, G show gross carboxylation rate equation (R_C_); B, E, H show carboxylation rate when the canonical PGS pathway is included; and C, F, I show the carboxylation rate when the alternate PGS pathway is included at 0, 25, and 55°C, respectively. A–F share a color scale; G–I have a different color scale.

Decreasing temperature reduced the difference in carboxylation speed between forms IAc and II. At 0°C, the two enzymes had at most a 2 carbon per second (C/s) difference (Fig. 3A–C). At 25°C, form II was up to 11 C/s faster than IAc (Fig. 3D–F), although form IAc outcompeted form II over a larger CO_2_/O_2_ parameter space. Both of these trends continued at 55°C, where form II is up to 55 C/s faster (Fig. 3G–I).

### Thermal adaptation analysis

We tested whether the Rubisco genes within our subzero genomes encoded a signal of thermal adaptation at the amino acid level. Applying the method of Raymond-Bouchard et al. 2018 [50], we scored a signal for hot or cold adaptation based on over 10 indices for each Rubisco (form II, IAc, and IAq) against a mesophilic average (see Methods for details). Of the 44 Rubisco sequences present in the genomes, 19 (43%) displayed a signal of thermal adaptation (Fig. 2B). Of the cold-adapted sequences, most were Rubisco form II sequences and included all of the clade 2 form IIs (those from subzero environments). The other cold-adapted form II sequences were found in cultured isolates that were able to grow at ≤5°C (Fig. S8, Table S1), possibly indicating cold adaptation. All form IIs with a signal of cold adaptation had similar amino acid substitutions and gaps (Fig. 4, Fig. S9). Form II in the type strain of *T. arctica* had the highest number of cold-adapted indices of the group, with 7 of the 10 indices pointing towards cold adaptation.

**Fig. 4:**
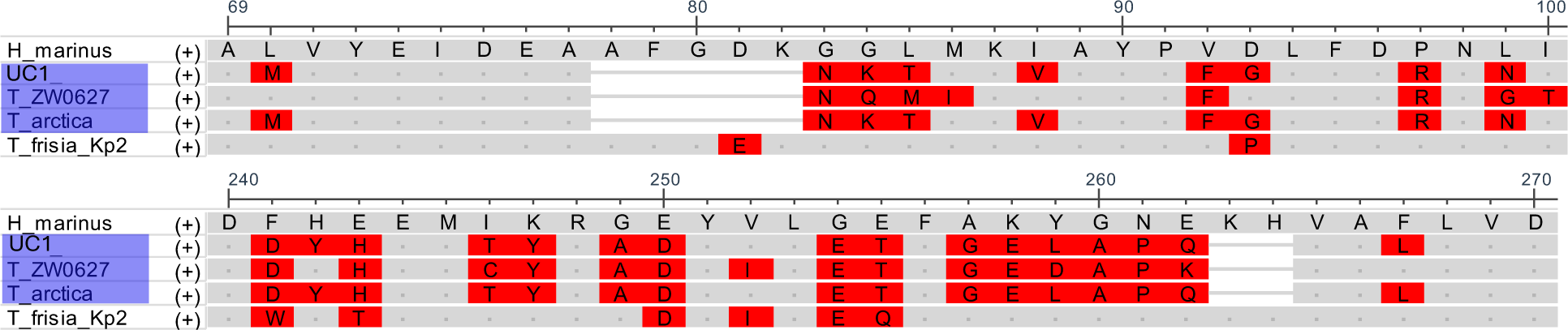
Partial amino acid alignments of Rubisco form II. Rubisco form II sequence excerpts from *H. marinus*, *T. frisia* Kp2, *T. arctica*, *T. arctica* UC1, and *T.* sp. ZW0627 around common, prominent substitutions/gaps found within the <cold-adapted= sequences (blue) compared to the mesophilic *H. marinus* and *T. frisia* Kp2. Amino acids highlighted in red show deviation from the *H. marinus* sequence.

Form IAc was the only Rubisco form with indications of hot adaptation. In particular, genomes with only form IAc all showed a signal of hot adaptation in that gene. Every MAG in Clade 3, which were all sequenced from hydrothermal vent fields, fell into this category. No form IAq sequence was cold- or hot-adapted.

### Rubisco form II structural analysis

To investigate potential structural differences in our “cold-adapted” form II Rubisco, a protein structure of the *T. arctica* UC1 form II sequence was compared to a reference mesophilic form II from *H. marinus* (Fig. 5). Proteins were created as dimers using AlphaFold [51,52]. There was no difference in the active site residues (Fig. S10A). However, many amino acid substitutions or gaps in the UC1 form II were found on the surface, primarily near the dimer-dimer interface (Fig. 5C). The gaps present in UC1 form II corresponded to an absence of residues that protrude from the surface of the protein in *H. marinus* (amino acids 78–82, Fig. 5B). In addition, the active site was more exposed in UC1 form II because of two substitutions (Y304P and P312A, Fig. 5A). These differences were not present in a comparison of the mesophilic *T. frisia* Kp2 form II to the *H. marinus* form II (Fig. S10B–C).

**Fig. 5:**
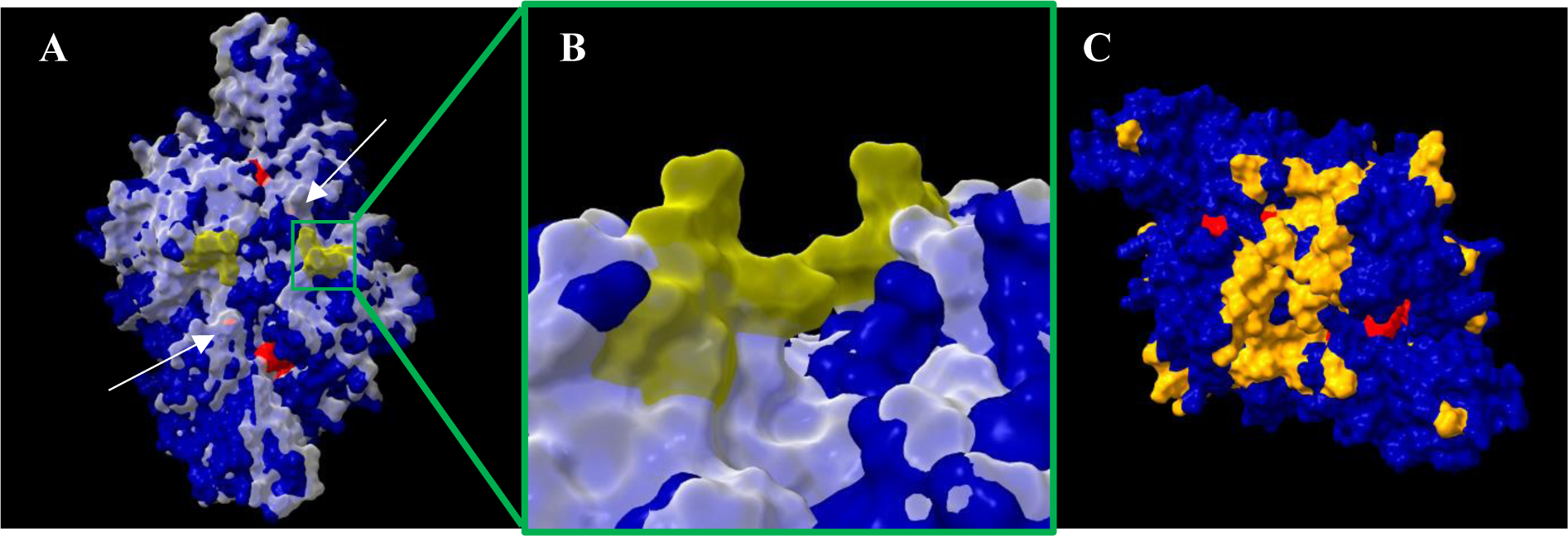
Rubisco form II structural differences between *T. arctica* UC1 and *H. marinus.* A) AlphaFold-generated [51,52] structures for *H. marinus* (transparent white) and *T. arctica* UC1 RbcII (dark blue) dimers aligned with ChimeraX [53]. Active sites colored red. Arrows point to the additional exposed active site residues in UC1 compared to *H. marinus*. The missing residues in UC1 at amino acid positions 78–82 (Fig. 4) are colored yellow on *H. marinus* structure (close-up view in B). C) UC1 form II dimer with residue differences to *H. marinus* that confer a semi-conservative or non-conservative change highlighted in orange.

## Discussion

### The CBB cycle in subzero environments

Subzero habitats on Earth represent a unique and rapidly declining environment, and provide an analogue for astrobiologically-relevant bodies like Mars, Enceladus, and Europa. In every subzero environment where we found the potential for chemolithoautotrophy, the CBB cycle was present, sometimes as the only known autotrophic pathway. What was particularly striking was that the absence of oxygen did not affect the presence of the CBB cycle. The CBB cycle is less thermodynamically favored under anoxic conditions given its higher energy requirements as compared to anaerobic carbon fixation pathways like the Wood-Ljungdahl pathway or the rTCA cycle [5] and thus is likely to be outcompeted. And yet it was present even in anoxic environments (Lake Vida brine [33] and cryopeg brines).

Our survey was conservative, as we only considered a taxon to be autotrophic if the study explicitly mentioned autotrophy or if reported taxa were known to always encode an autotrophic pathway. This approach may have biased our data towards the CBB cycle as it is better studied than the other carbon fixation pathways. For example, Thaumarchaeota were reported at several of the sites considered. Ammonia-oxidizing Thaumarchaeota use a modified 3-HP/4-HB cycle to fix carbon, but we cannot state conclusively that these detections equate to the potential for autotrophy as heterotrophic Thaumarchaeota were recently discovered [54,55]. A deeper re-analysis of past studies using available metagenomes and 16S sequences could reveal more autotrophy in subzero environments.

Despite these caveats, our survey demonstrates the pervasiveness of the CBB cycle at subzero temperatures even in oxygen-deprived habitats. While the CBB cycle is relatively understudied in chemolithoautotrophs and anaerobes, it could hold signals for diversification across widespread environments that would be relevant to the search for autotrophy beyond Earth.

### Rubisco forms: driven by taxonomy or suitability for the environment?

Rubisco catalyzes the key step of the CBB cycle and exerts an important biosignature on carbon isotopes that is used as evidence for life [56]. Rubisco forms are distinct between taxonomic groups but also possess different kinetic traits [12]. Most research on Rubisco has focused on the form IB found in plants, which (with a few exceptions) is only able to photosynthesize above 0°C; it is generally thought that form IB Rubisco is functionally inactive at subzero temperatures [57]. Exploring the different Rubisco forms in subzero environments has provided insight into both the taxonomic composition of the autotrophic community and which Rubisco forms may be functional at subzero temperatures.

We found a diverse range of bacterial Rubisco forms in our samples, including II, IA, IC, and IE. While forms II and IA were found in both cryopeg brines and sea-ice samples, form IC was only found in sea-ice samples, and form IE was only found in cryopeg brines. Form II and form IA are commonly found in proteobacteria, which formed a high percentage of the community in both environments [37]. Form II, however, was several times as abundant than form IA in both our cryopeg brines and sea-ice samples. Form II is typically considered to be faster than form I but less specific for CO_2_ [48], which would make it more beneficial than IA in an anoxic or low-oxygen environment, such as the cryopeg brines. On the other hand, sea ice is oxygenated, so form II’s prevalence in the sea-ice samples may instead be due to the increased solubility of CO_2_ versus O_2_ at low temperatures. It is also possible that form II is simply more common than form IA in all proteobacteria, and thus in our subzero population. We do not have enough information about the prevalence of either of these forms in proteobacteria to differentiate between these possibilities.

Form IE was recently discovered in soil and atmospheric Actinobacteria [12] and is known to occur in high Arctic deserts [58]. Many of the IE sequences were taxonomically assigned to the genus *Microbacterium*, which is a common member of permafrost communities [59]. Thus, the presence of form IE in one of our cryopeg brine MAGs may reflect the terrestrial influence of the surrounding permafrost on the cryopeg brines. Form IC is found in proteobacteria, which are ubiquitous, and we expected to find it in both sea-ice samples and cryopeg brines. Its absence in the cryopeg brines could be due to a population bottleneck after cryopeg formation [60] or to a kinetic disadvantage in the cryopeg brines versus sea ice. Current measurements of form IC kinetics are scarce but suggest that it is comparable to form IA [8]. Kinetics have not been measured for form IE, so a direct comparison of these two forms is not possible.

Both sea ice and cryopeg brines support a range of microenvironments [36]. Because of the variability of the kinetics of different Rubisco forms, the presence of multiple forms of Rubisco in one community could maximize the amount of carbon fixation the community could perform in a variable environment. Thus, the wide variety of Rubisco forms found in these environments may be a useful advantage for these communities.

### Thiomicrorhabdus as a model organism to study multiple Rubisco forms

One particular genus, *Thiomicrorhabdus*, was detected in subzero environments across both poles. Members of the genus were measured actively fixing carbon in Lost Hammer Spring and Gypsum Hill Spring in the Canadian Arctic [30,41] and in the outflow brine from Blood Falls in Antarctica [29].

The prevalence of *Thiomicrorhabdus* in subzero environments presented a unique approach to study Rubisco form and function across extreme environments. The genus is found in many different habitats including hydrothermal vents, coastal sediments, and brines [38]. While cultured strains have been reported to grow from –2 to 45°C, they have been detected in environments with temperatures as low as –13°C [33].

The majority of *Thiomicrorhabdus* genomes have three different forms of Rubisco (II, IAc, IAq) which are also found in the *Hydrogenovibrio* outgroup, suggesting the common *Thiomicrorhabdus* ancestor contained all three forms. Laboratory cultures of *H. marinus* found differential expression of the three forms under different CO_2_ concentrations [61], with form II expressed under low O_2_, and form IAc expressed with its carboxysome under low CO_2_. The wide range of temperature, pressure, and salinity across *Thiomicrorhabdus* habitats also influences CO_2_ and O_2_ solubility. Sediments, where many *Thiomicrorhabdus* species reside, support a range of microenvironments with varying O_2_ and CO_2_ conditions and thus retention of multiple Rubisco forms would be advantageous.

However, there are multiple genomes where one or two Rubisco forms have been lost, and this pattern cannot be explained by a few loss events. This winnowing of Rubisco forms present in a species may represent a specialization to a narrower range of conditions. Clade 2 of *Thiomicrorhabdus* is a good candidate for this type of loss, as all of the clade’s species are present in permanently cold environments where CO_2_ solubility would be higher and a carboxysome would be less needed. The loss of Rubisco genes could also be due to increased evolutionary pressure on the species such that genome streamlining occurred, if duplicate Rubisco genes did not provide enough advantage to offset the nutritional cost of keeping them in the genome.

### Rubisco adaptation to extreme environments

Form II is likely to be important at subzero temperatures. It is the only form found in all of the genomes isolated from subzero environments and has a strong signal of cold adaptation that correlates with subzero habitats or a low minimum growth temperature. A preference for form II at low temperatures is consistent with its known faster kinetics than form I [48]. For our carboxylation model, we used form II from the closely related *Hydrogenovibrio*, which has the third-fastest form II measured to date [48] and in our model clearly outcompetes form I over most of the parameter space, including at cold temperatures. Interestingly, cold temperatures lowered the competitive advantage of form II. However, our modeling was dependent on extrapolating kinetic properties from only a limited dataset, which was particularly sparse for temperature sensitivity. Also, our finding of cold adaptation within the form II protein sequence suggests that the kinetics of form II from subzero environments may display different kinetics to what we modelled.

To date, there is little evidence of cold adaptation of Rubisco. Studies of form I Rubisco kinetics from C3 plants from cool climates [62] and from psychrophilic algae [63,64] have shown little evidence of cold adaptation. However, some cold-water seaweeds show evidence of faster Rubisco kinetics at cold temperatures [21], and there is some evidence that the psychrophilic diatom *Fragilariospsis cylindrus* has slightly faster carboxylation at cold temperatures (<10°C) [65].

In general, Rubisco is under strong purifying selection [14]. A previous study of 7 chemoautotrophic Piscirickettsiaceae family genomes (including *H. marinus*) found that all three forms of Rubisco had evolved under purifying selection, especially forms IAc and II [66]. However, a growing number of studies [14,67] have highlighted signals of adaptation within a few key amino acids that can have a significant effect on kinetics. Expanding our knowledge of Rubisco functional diversity and kinetics could be exploited in the search for life in extreme environments.

Our study did not focus on Rubisco forms at high temperatures. Nonetheless, we did observe the importance of form IAc in hydrothermal vent MAGs, which also displayed a signal for high temperature adaptation in the protein sequence. Form IAc is associated with a carboxysome, which would be advantageous at high temperatures where CO_2_ solubility would be low. Rubisco proteins in thermophiles have a higher temperature optimum of activity [68–71], although accessory proteins like the Rubisco Activase are typically considered more important for the activity of Rubisco at high temperatures than the thermostability of the Rubisco protein itself [57].

The benefit of retaining IAq is more difficult to discern. Our modeling found no condition in which form IAq outcompeted form II or IAc, and we found no evidence of thermal adaptation in any sequence. A previous study of *H. marinus* also found no CO_2_ condition where form IAq was the predominant Rubisco expressed [61]. We speculate that form IAq may be useful when resources are too low to synthesize a carboxysome, which requires a high amount of nutrient investment to build the protein shell, but O_2_ levels are too high for form II to function effectively.

We did not investigate potential adaptations to salinity in these Rubisco sequences, despite the high salinities of many of the environments studied in this work. Based on a theorized incompatibility between the types of protein changes needed for hypersaline adaptation and psychrophilic adaptation [72], the evidence of cold adaptation in the form II Rubisco sequences may preclude a simultaneous adaptation to high salt conditions. Despite theoretical contentions, extracellular enzymes produced by psychrophilic bacteria have been shown to be active, in some cases preferentially active, under combined subzero and hypersaline conditions [73]. The environmental pressure to adapt to such extreme conditions would be limited for intracellular proteins, as they are not subjected to the ionic concentrations present for extracellular proteins, particularly in organisms that use compatible solutes, a common form of salt protection [72]. Regardless of potential adaptation in the Rubisco sequences themselves, salinity does affect the availability of CO_2_ and O_2_. High salinity environments have lower gas solubility and therefore lower CO_2_ and O_2_ concentrations, potentially increasing the need for a carboxysome to provide sufficient CO_2_ to Rubisco.

### Comparing different Rubisco forms by modeling kinetics

We developed a modeling framework to compare carboxylation rates of different Rubisco forms to test under which environmental niches Rubisco forms would outcompete each other, beyond the canonical atmospheric conditions at 25°C so often quoted. This approach allowed us to explore the environmental limits of autotrophy and could be used to identify viable environmental conditions for autotrophy on other worlds (e.g., in Martian subsurface brines or the Enceladus ocean).

A unique component of our model is the incorporation of the carbon loss due to PGS pathways resulting from the Rubisco oxygenase reaction. There are four known PGS pathways with different carbon costs [74], and it is not known which pathway is utilized in *Thiomicrorhabdus*. The C2 cycle (e.g., in plants) and the glycerate pathway both cost one carbon for every two phosphoglycolate molecules processed, whereas the malate and oxaloacetate pathways cost two carbons for each phosphoglycolate processed [74]. Only when a PGS pathway is added to the model is there a parameter space where form IA outcompetes form II, and also where Rubisco carboxylation is no longer a viable strategy for autotrophy. Outside of *Thiomicrorhabdus*, this same approach can be applied to other Rubisco proteins with known kinetics in other species, and to approximate specific environments of interest. We found that under low oxygen conditions, as in a marine oxygen deficient zone, form II is likely to be more common, and may be a better model for carbon fixation on Enceladus or Europa, which are thought to be anoxic [75]. In contrast, Earth’s current atmospheric conditions of high O_2_ and low CO_2_ (for geologic time) result in both form II and IA losing carbon; in these cases, a carboxysome might be utilized to further exclude oxygen and increase CO_2_ around Rubisco. The current CO_2_ and O_2_ regime also supports the dominance of form IB in the modern Earth atmosphere, as form IB in plants has a higher S_C/O_ than form IA [10].

## Conclusion

Our study found multiple lines of evidence that form II Rubisco has a competitive advantage at cold temperatures. First, it is the most abundant autotrophic bacterial form found in our metagenomes from subzero cryopeg brines and sea ice. Second, it is the only form retained in all *Thiomicrorhabdus* genomes from cold environments. Third, our model indicates it performs better than Rubisco form I over the majority of conditions at cold temperatures. Fourth, many of the form II Rubisco amino acid sequences contained a signal of cold adaptation and have potential structural changes on the surface of the protein.

Our study also shows that the use of models that incorporate PGS, and the exploration of multiple forms of Rubisco within a single genus, can provide valuable information to probe the environmental limits at which Rubisco and the CBB cycle can operate. Ultimately, this work underscores the need for more data on Rubisco kinetics from non-plant organisms, especially chemolithoautotrophic bacteria, and the importance of studying autotrophy in extreme environments for understanding current life on Earth and potential life on other planetary bodies.

## Methods

### Autotrophy at subzero

The prevalence of different autotrophic pathways in subzero environments was determined by a survey of primary literature. If autotrophy was mentioned, autotrophic members were categorized and noted. If autotrophy was not mentioned, community members from genera or families known to be obligate autotrophs were considered to be autotrophic and listed with their known autotrophic pathway. Only chemolithoautotrophs were included for this analysis. If available, temperature information from the same study was used. If not, temperature was collected from other studies of the same habitat.

### Identification of Rubisco genes from cryopeg brines and sea ice

Rubisco genes were identified from metagenomes from cryopeg and sea-ice sackhole brines and sea-ice top and bottom core melts in May 2017 and May 2018, sampled as described in [36]. Briefly, sea-ice from ∼1.1-1.2 m thick first-year land-fast sea ice were sampled ∼100m from the Barrow Sea Ice Mass Balance site (71.37219° N, 156.53828° W) in early spring [36]. Sea-ice brines were collected from sackholes, and sea-ice core melts (from the top third and bottom third of the core) were collected from the same ice field. Cryopeg brines were sampled from the Barrow Permafrost Tunnel (71.2944° N, 156.7153° W), 6 meters below the surface. Brines were extracted from approximately 2 meters below the tunnel floor using a hand pump and sterile receiving system. Sample nomenclature and environmental data are in Table S6. Further site and sampling details can be found in [36,37].

From these samples, a set of 11 metagenomes was previously generated and described by Rapp et al. 2021 [36], Briefly, sequencing was done at the DOE Joint Genome Institute (JGI) on an Illumina NovaSeq sequencer, and data from individual samples was subsequently quality-controlled, assembled, structurally and functionally annotated through the DOE–JGI Metagenome Annotation Pipeline (MAP v4.16) [76]. Coverage information was obtained by mapping filtered reads back to assembled contigs using BBMap v38.25 with “ambiguous = random”. Gene taxonomy was inferred through a homology search using LAST [77] against IMG reference isolates (high quality public genomes), and scaffold lineage affiliation then represents the last common ancestor of all best gene hits on a scaffold, provided that at least 30% of the genes had hits.

All nucleotide sequences annotated as the Rubisco large subunit were translated into protein sequences with any sequences less than 100 amino acids discarded. Amino acid sequences were aligned using MAFFT [78] with 66 reference sequences encompassing different Rubisco forms from Prywes et al. 2023 [12]. A maximum likelihood tree was built using IQTREE2 [42] and Rubisco form was assigned based on the closest reference sequences.

Forms ID and IC do not form separate clades and are typically assigned based on host (ID is used for eukaryote sequences, while IC is used for prokaryote sequences) [12]. Thus, all ID and IC sequences were run through NCBI’s BLAST tool [79] in order to confirm that ID sequences matched known eukaryotic sequences while IC sequences matched known prokaryotic sequences. If not, they were re-assigned manually.

Pie charts were created based on the coverage of bacterial Rubisco sequences separated by form in all cryopeg brine samples and all sea-ice samples, using the abbreviations from [12]. All form IV subtypes were counted as one form IV category in the charts, as form IV does not participate in autotrophy.

### MAG Assembly

For the reconstruction of metagenome-assembled genomes (MAGs), quality-controlled paired-end Illumina reads of cryopeg brine samples CBIW_0.2_2017, CBIW_3.0_2017, CBIW_2018, CBIA_2018 and CB4_2018 were co-assembled using MEGAHIT v1.2.1 [80]. The resulting cryopeg co-assembly was processed with ‘anvi-script-reformat-fasta’ for input into Anvi’o v6.2 [43], retaining only contigs of 2,500 bp and longer. The quality-controlled input reads of each sample were mapped back to the co-assembly using Bowtie2 v2.3.0 [81] to generate sequence alignment map files, which were then converted to binary format using SAMtools v1.5 [82], and sorted and indexed using ‘anvi-init-bam’ [43].

The co-assembled metagenome contigs were binned using MetaBAT [83], MetaBAT2 v2.15 [84], MaxBin2 v2.2.7 [85], BinSanity v0.4.0 [86] and GroopM2 v0.2.1 [87], and the resulting bin sets were integrated into one combined draft ensemble set using DAS Tool [88], metaWRAP [89] and uniteM (https://github.com/dparks1134/UniteM). The draft ensemble was optimized and dereplicated using another iteration of DAS Tool, metaWRAP and uniteM to produce three sets of non-redundant draft bins. Bin completion and contamination was determined using CheckM v1.1.3 [90], and only the highest quality draft bin set was retained. The retained bins were then imported into Anvi’o for vizualisation and manual curation based on sequence composition and differential coverage [43], and the quality and completion of the refined bins was assessed with CheckM v1.1.3 [90]. Only bins of 50-100% completeness and 0-10% redundancy (potential contamination) were retained, corresponding to medium-, high-quality and complete MAGs [91].

### MAG Analysis

MAGs were classified taxonomically using the classify_wf pipeline of GTDB-Tk v0.3.2 with database release r89 [92]. Prodigal v2.6.3 [93] was used for gene calling as implemented in Anvi’o [43], KofamKOALA v2020-07-01 with KEGG release 95.0 [94] was used to annotate functions with KEGG Orthologs (KO), and KEGG decoder v1.2 [95] was used to contextualize KO assignments within metabolic pathways and to assess pathway completion.

Estimates of the relative contribution of MAGs to the total community composition in individual cryopeg brines was calculated based on read mapping to MAGs using CheckM v1.1.3 [90] with the “profile” command. An Average Nucleotide Identity (ANI) pairwise comparison using fastANI [96] was done to confirm genus identity and determine species identity. ANI values >95% were considered to indicate a shared species identity.

### Thiomicrorhabdus phylogenetics

A total of 45 additional *Thiomicrorhabdus* genomes, including 28 MAGs and 17 isolate genomes, and 2 isolate genomes from a closely related genus, *Hydrogenovibrio*, were obtained from the NCBI database (Table S1). Each genome was analyzed in CheckM [90] using pplacer [97], prodigal [98], and HMMER [99], for completeness and contamination using as markers the 638 single-copy genes common to the Pisciricketsiaceae family, to which the *Hydrogenovibrio* and *Thiomicrorhabdus* genera both belong. Genomes under 70% completeness or above 5% contamination were discarded. Genomes were also compared pairwise for ANI using fastANI [96]. Genomes matching at ≥95% ANI were de-replicated, and the most complete representative was chosen to represent the cluster for phylogenetic analysis. Two genomes that otherwise met these criteria were excluded from further analysis for lack of any complete Rubisco genes.

Functional annotation of the remaining 24 genomes was done using the Snakemake [100] “anvi-run-workflow -w contigs” workflow in Anvi’o [43] against the KEGG [101–103], COG [104,105], and Pfam [106,107] databases. Environmental data and location of sample were collected from original sources (Table S1). A map of sample locations was created using QGIS 3.30.1 [108] and the ESRI Ocean Basemap [109].

A phylogenetic tree of the *Thiomicrorhabdus* genomes was constructed using the 71 phylogenetically informative marker genes of the Bacteria_71 collection in Anvi’o [43–45]. For each species, nucleotide sequences annotated as these marker genes were extracted from the genome and translated into amino acid sequences using Prodigal [98]. In cases where redundant markers were recovered for the same species, the gene with the longest sequence was selected. For each marker, predicted protein sequences were aligned individually using mafft [78] and alignments were subsequently trimmed using trimal (-gt 0.1) [110]. Next, individual marker alignments were concatenated into a per-species supermatrix. Only those species with 35 or more of the 71 marker genes were retained. Finally, a maximum-likelihood species tree was inferred using IQTree2 with a model selection step and 1,000 ultrafast bootstraps (-bnni -m TEST -st AA -bb 1000) [42]. The resulting topology was then visualized in iToL [46] using *Hydrogenovibrio marinus* and *H. thermophila* as an outgroup.

### Thiomicrorhabdus Rubisco analysis

For each *Thiomicrorhabdus* and *Hydrogenovibrio* genome, all genes annotated as Rubisco large subunits (RbcL) were aligned and trimmed. RbcL genes clustered into three groups which corresponded to the three known forms (form II, form IAc, and form IAq) from *H. marinus* [61]. A manual confirmation of this assignment was performed based on species with known Rubisco forms from [47]. Small subunits were identified based on their proximity to large subunits with an additional manual check to confirm that all form I sequences had a small subunit and a large subunit, and that no form II sequence had a small subunit.

Genome annotations were used to create Rubisco cassette gene maps using ggplot [111] and gggenes [112]. A LysR gene next to an RbcL or RbcII gene was considered the beginning of a RbcII or RbcIAq cassette respectively, while a CynT or CbbO gene was considered the ending of same. RbcIAc cassettes were considered to start at a RbcLIAc gene and end at a LysR gene. Occasionally gene rearrangements or contig ends/starts necessitated adjustments to the start or end of the cassette depiction.

### Theoretical model

The carboxylation reaction rate of a Rubisco enzyme (R_C_) is competitively inhibited by O_2_ and can be modeled under different CO_2_ and O_2_ concentrations using a competitive-inhibitor Michaelis Menten equation as described in Flamholz et al. 2019 [8] (Eqn. 1):

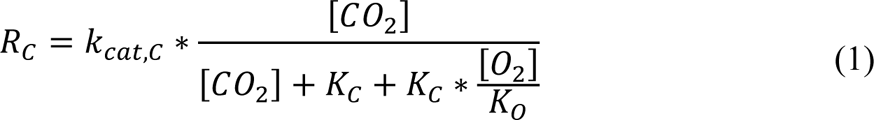

where R_C_ is the gross carboxylation rate of Rubisco, *k*_cat,C_ is the carboxylation rate of each active site (CO_2_/s), K_C_ and K_O_ are the half saturation constants for CO_2_ and O_2_, respectively, and [CO_2_] and [O_2_] are the concentrations of CO_2_ and O_2_, respectively. This equation can be used to directly compare Rubisco carboxylation of different enzyme forms.

To include the carbon loss due to the PGS pathway, which is the result of the oxygenation reaction, we need to calculate the oxygenation reaction rate (R_O_) using a similar equation (Eqn. 2):

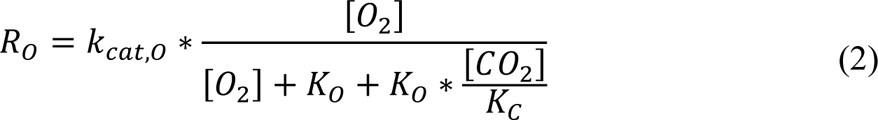

The value for *k*_cat,O_ is not measured directly; instead, we utilized the commonly measured specificity factor (S_C/O_), which is the ratio of the carboxylase/oxygenase reaction (Eqn. 3):

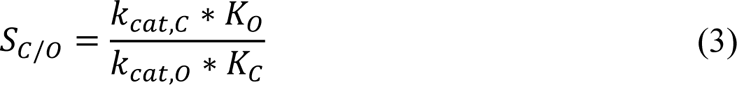

The relationship between S_C/O_ and R_C_/R_O_ can be described by Equation 4, which can be rearranged to calculate R_O_.

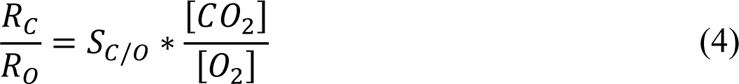

While Flamholz et al. 2019 [8] argue that this relationship only exists during low CO_2_ and O_2_ conditions, when CO_2_ << K_C_ and O_2_ << K_O_, we find, in agreement with the original derivation by Laing et al. 1974 [113], that this relationship is true for any CO_2_ or O_2_ concentration (see supplemental Methods for full derivation).

The canonical salvage pathway (the C_2_ cycle) for Rubisco’s oxygenated product, 2-phosphoglycolate (2-PG), results in a loss of 1 CO_2_ molecule for every 2 2-PG molecules fixed [59]. Using this stoichiometry, the net carboxylation rate (*f*) is:

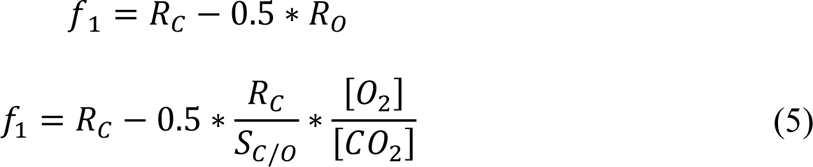

Three other salvage pathways for 2-PG have been found in bacteria [74]. The glycerate pathway has the same stoichiometry as the C_2_ cycle, but the malate cycle and the oxalyl-CoA decarboxylation route both result in a loss of 2 CO_2_ molecules for every 2-PG molecule fixed. The net carboxylation rate using this stoichiometry (called the alternate net carboxylation rate) is:

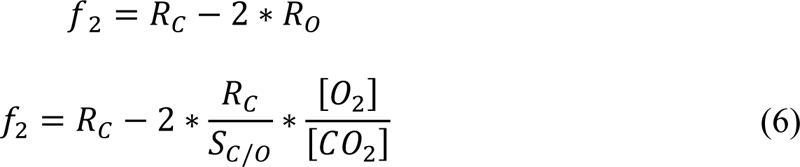

A model for each of the possible tradeoffs (the gross Rubisco carboxylation rate, Rubisco carboxylation – the canonical PGS pathway, and Rubisco carboxylation – the alternate PGS pathway) was created for each Rubisco. A rate of 0 or less denoted that autotrophy could not occur under the specified conditions.

To include temperature effects in the model as well as the effect of CO_2_ and O_2_ concentrations, an Arrhenius-type equation with the form 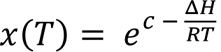 was created for each kinetic parameter (*k*_cat,C_, K_C_, K_O_, and *k*_cat,O_) for each enzyme, as in Galmes et al. 2016 [19]. For each equation, an appropriate ΔH was taken from reported values in Galmes et al. 2016, while the constant *c* was calculated by setting the equation equal to a reported value for each kinetic parameter. *R* is the universal gas constant, 0.00831 kJ/mol/K.

The following shows an example calculation for the *k*_cat,C_ for form IAc: The reported value for the *H. marinus* form IAc *k*_cat,C_ is 2.98 at 30°C [49], and the ΔH chosen for form IA *k*_cat,C_ is 47.2 kJ/mol (an average of three proteobacteria values) [11]. Thus:

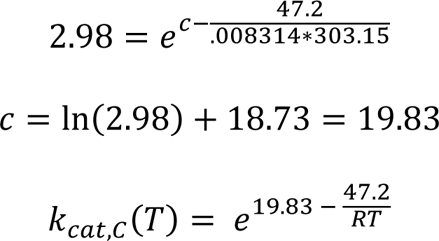

Values used for this analysis are available in Tables S3 and S4, but briefly: reported kinetic parameters were taken from Hayashi et al. 1998 [49], Davidi et al. 2020 [48], and Flamholz et al. 2019 [8] (Table S4); ΔH values were taken from Galmés et al. 2016 [19] and Galmés et al. 2015 [11] (Table S3). As the studies that reported ΔH values for form IA enzymes did not distinguish between form IAc and form IAq enzymes, the form IAc and form IAq models use the same ΔH values.

To compare Rubisco forms, the rate of the form IAc Rubisco was subtracted from form II. When the resulting equation has a positive value, form II is faster than form IAc, and vice versa.

### Rubisco thermal adaptation

There are a number of amino acid substitutions that are associated with either cold or hot adaptation [114], and while these may suggest possible thermal adaptation, they do not guarantee it [50]. Thus, amino acid sequences of Rubisco were interrogated for signals of potential thermal adaptation using the methods from Raymond-Bouchard et al. 2018 [50]. Ten indices were incorporated to calculate an adaptive score: number of glycines; number of prolines; number of serines; number of acidic residues (D,E); number of charged residues (D,E,H,K,R); number of polar uncharged residues (C,N,Q,S,T,Y); aliphatic index; aromaticity; and hydrophobicity. Rubisco sequences included the large and small subunits, where relevant, and counts of all amino acids, as well as aromaticity, hydrophobicity (using Grand Average of Hydropathicity index, hereafter GRAVY), and isoelectric point were calculated using the Biopython [115] module Prot.Param, modeled on the Expasy ProtParam tool [116] for each sequence. Aliphatic index was calculated as in Ikai 1980 [117].

A reference (mesophilic) value for each index, within each Rubisco form (IAq, IAc, and II), was determined as the average and standard error value of Rubiscos from five genomes *(H. marinus, T. cannonii, T. frisia* Kp2*, T. heinhorstiae,* and *T. sediminis*) based on their known mesophilic temperature ranges. Only four Rubisco sequences were used to generate the reference values for form IAq, as *T. sediminis* does not contain a form IAq Rubisco. A one-sample t-test was used to determine if the test sequence index value was significantly different from the mesophilic reference values. Cold or hot adaptation was assigned to the significant indices based on direction of change (Table S7).

An index with a significant value in the cold direction scored as +1, and in the hot direction as –1. Those with no significance scored 0. The total score of all 10 indices were summed to receive an overall thermal adaptation score. Values between –2 and 2 were considered neutral. Values ≥3 indicated cold adaptation, while values ≤ –3 indicated hot adaptation.

### Rubisco 3D structure

Sequences of *H. marinus* form II Rubisco, *T. frisia* Kp2 form II, and UC1 form II proteins were used to build hypothetical Rubisco dimer protein structures using the AlphaFold [51,52] multimer model. Resulting PDB files were visualized in ChimeraX [53] and aligned in space using the in-built alignment tool (only paired atoms). Amino acids associated with the active sites were highlighted in red for each protein. Amino acids in *H. marinus* missing in UC1 and all other “cold” Rubisco form II were highlighted in yellow. In addition, amino acid changes in UC1 where the function of the amino acid changed significantly, i.e., where the BLOSUM62 matrix identifies the change as being semi-conservative or not conservative (consensus symbol “.” or empty on the CLUSTAL description), were highlighted in orange.

## Data Availability

Metagenomic data used for the identification and extraction of Rubisco sequences from cryopeg brines and sea-ice samples are available under SRA Bioproject accession numbers PRJNA570544, PRJNA539724, PRJNA539309, PRJNA539308, PRJNA539600, PRJNA539601, PRJNA518292, PRJNA518293, PRJNA570545, PRJNA518295 and PRJNA518294. Cryopeg brine metagenomes used for the generation of MAGs are available at PRJNA539724, PRJNA539309, PRJNA539308, PRJNA539600, PRJNA539601. Autotrophic cryopeg brine MAGs were deposited in TBD (attached as fasta files currently, but will be available before publication). All other publicly available genome sequences used for this study can be found in NCBI under the accession numbers listed in Table S1.

## Author Contributions

K.H. and J.Y. designed and wrote the original manuscript. All authors contributed to edits and revisions. K.H. and J.Y. performed the primary literature search. J.Z.R. performed the metagenomic analyses of cryopeg brines and sea-ice samples and extracted the Rubisco sequences. A.L.J. performed the *Thiomicrorhabdus* genus phylogenetic analysis and provided assistance and advice on phylogeny and alignment. K.H. performed all other work.

## Acknowledgements

We would like to thank Kathleen Scott for her discussions with us on *Thiomicrorhabdus*, and Zachary S. Cooper for his input on the cryopeg brine environment. This work was funded by the Simons Foundation Award 561645, the Sloan Fellowship, and NSF CAREER Award 2142491. J.Z.R. and J.W.D. were supported by the Gordon and Betty Moore Foundation grant no. 5488 and the Karl M Banse Professorship (JWD). A.L.J. was funded by the Stanford Science Fellows program and the NSF Postdoctoral Fellowship in Ocean Sciences.

